# The microstructure-weighted human connectome: network properties and structure-function correlations across spatial scales

**DOI:** 10.64898/2026.05.19.726180

**Authors:** Arthur P C Spencer, Saina Asadi, Yasser Alemán-Gómez, Qiaochu Wang, Maciej Jedynak, Chun Hei Michael Chan, Alexandre Cionca, Dimitri Van De Ville, Olivier David, Patric Hagmann, Ileana Jelescu

**Affiliations:** Department of Radiology, Lausanne University Hospital (CHUV), Lausanne, Switzerland; Faculty of Biology and Medicine, University of Lausanne, Lausanne, Switzerland; Neuro-X Institute, École Polytechnique Fédérale de Lausanne (EPFL), Geneva, Switzerland; Institut de Neuroscience des Systèmes, Aix Marseille University, Marseille, France; Department of Radiology and Medical Informatics, University of Geneva, Geneva, Switzerland; Department of Neurosurgery, Fondation Lenval Pediatric Hospital, Nice, France

**Keywords:** Connectomics, biophysical diffusion modelling, diffusion MRI, functional MRI, structure-function coupling, SEEG

## Abstract

Conventional connectome edge weights, such as number of streamlines (NOS) or diffusion tensor imaging (DTI) metrics, lack specificity to microstructural details which may hold relevance for macroscale brain organisation. Since biophysical diffusion modelling offers greater specificity to microstructure, we investigated whether parameters from the Standard Model of white matter diffusion provide informative alternatives for connectome weights – namely the intra-axonal signal fraction (f) and perpendicular extra-axonal diffusivity (D_e_ ^⟂^). Using diffusion MRI data from healthy adults, we constructed structural networks at four parcellation scales, weighted by f, D_e_ ^⟂^, NOS, fractional anisotropy (FA) and radial diffusivity (RD). While all weights reproduced expected small-world properties and hub structure, only D_e_ ^⟂^ and normalised NOS captured non-random properties of local organisation at all spatial scales. We then correlated each weighted connectome with resting-state fMRI functional connectivity and intracranial measurements of conduction velocity. At both the whole-brain and regional level, NOS gave strong coupling with fMRI functional connectivity. Connectomes weighted by D_e_ ^⟂^, FA and RD gave weak but significant coupling with fMRI functional connectivity and conduction velocity. Notably, D_e_^⟂^ and RD gave highest consistency in regional structure-function coupling across spatial scales. Thus, connectome weights derived from D_e_ ^⟂^ capture meaningful aspects of brain network organisation with functional relevance.

## 1 Introduction

White matter fibres provide the structural substrate which facilitates functional brain dynamics, mediating the transmission of information between grey matter regions (Filley & Fields, 2016). Thus, better characterising white matter connectivity may offer insight into functional interactions across the brain, including how these interactions are disrupted by disease or disorder (Zalesky & Di Biase, 2026). In network representations of the brain, each edge can be assigned a weight which represents the strength of the white matter connection between a pair of grey matter regions, enabling mathematical characterisation of brain organisation using graph-theoretical analysis (Bassett & Sporns, 2017; Bullmore & Bassett, 2011; Fornito et al., 2013; Hagmann et al., 2010). Traditionally, number of streamlines (NOS) is used as a connectome edge weight, but this is highly dependent on tractography parameters and normalisation strategies (Calamante, 2019; Jones et al., 2013). Therefore, this does not provide an objective quantity with direct relevance to the underlying fibres. Alternatively, many studies have used diffusion tensor imaging (DTI) metrics, such as the mean fractional anisotropy (FA) along streamlines, as a measure of connectivity (van den Heuvel & Sporns, 2011). However, while FA is sensitive to microstructure, it is not a specific measure of any given microstructural property (Jelescu et al., 2016b, 2020) and is strongly influenced by the presence of crossing fibres (Tournier et al., 2011). The efficiency of functional signal propagation along white matter tracts is dependent on microstructural properties including myelination and fibre density (Asadi et al., 2025; Caminiti et al., 2013; Drakesmith et al., 2019; Mancini et al., 2021). As such, parameters sensitive and specific to these microstructural features may provide connectome weights with maximal relevance to the functional organisation of the brain.

With advances in accelerated acquisitions, high angular resolution multi-shell sequences are now commonplace for diffusion-weighted imaging (DWI) studies (Dell’Acqua & Tournier, 2018; Jeurissen et al., 2014). The wealth of information available in such acquisitions can be used to probe microstructural information, using biophysical modelling to characterise diffusion properties with higher specificity to microstructural details (Jelescu et al., 2020; Jelescu & Budde, 2017). Despite this, many connectomics studies still use FA as connectome weights, thus employing only a subset of the information available in the DWI data. Instead, adopting microstructure parameters from biophysical modelling as connectome weights may offer additional detail about the macroscale organisation of brain networks.

Previous studies have demonstrated the utility of certain diffusion model parameters as connectome weights. Connectomes weighted by axon diameter distribution from axonal spectrum imaging (Gast et al., 2023) show relevance to cognitive performance (Gast & Assaf, 2024). Additionally, parameters from the neurite orientation dispersion and density imaging (NODDI) model (Zhang et al., 2012) provide connectome weights with sensitivity to age-related changes (Buchanan et al., 2020) and alterations in temporal lobe epilepsy (Lemkaddem et al., 2014). However, several studies have challenged the sensitivity of axon diameter estimations and the specificity of NODDI parameters (Howard et al., 2022; Huang et al., 2015; Jelescu et al., 2016a; 2015; Veraart et al., 2020). Further studies have explored myelin-sensitive measures as connectome weights, such as longitudinal relaxation rate (R1) (Boshkovski et al., 2021; Nelson et al., 2023) or g-ratio (Mancini et al., 2018), demonstrating their relevance for modular brain organisation and their sensitivity to changes induced by disease (Kamagata et al., 2019) or cognitive training (Caeyenberghs et al., 2016). However, these quantitative MRI measures require specific acquisitions which are not available in many retrospective datasets.

The Standard Model (Coelho et al., 2022; Liao et al., 2024; Novikov et al., 2018) is a biophysical model of water diffusion in white matter. This compartment model represents the diffusion signal as a collection of fascicles, each containing axons modelled as impermeable zero-radius cylinders, and extra-axonal diffusion modelled as an axially symmetric diffusion tensor. Fascicles within a voxel (assumed to have the same compartment fractions and diffusivities) are distributed according to a fibre orientation distribution function. The introduction of the Standard Model overcame the limitations imposed by oversimplified assumptions of previous biophysical models, providing parameters with greater specificity to microstructural details (Coronado-Leija et al., 2024; Jelescu et al., 2020; Jelescu & Budde, 2017). Notably in the context of white matter connectivity networks, the axon signal fraction (f) provides a measure of axon density, and the extra-axonal perpendicular diffusivity (D_e_^⟂^) is sensitive to myelination (Nguyen-Duc et al., 2026; Coronado-Leija et al., 2024; Jelescu et al., 2016b; Fieremans et al., 2012). While this increased biophysical realism comes at the cost of parameter degeneracies in certain cases, these degeneracies can be mitigated either through appropriate acquisition design (e.g. multi-TE acquisition) or fitting constraints (e.g. by excluding the free water compartment) (Coelho et al. 2024). The latter constraint enables the Standard Model to be readily applied to many conventional multi-shell DWI datasets.

In this study, we used Standard Model parameters f and D_e_ ^⟂^ as structural connectome weights, investigating network properties of these connectomes in comparison with networks weighted by DTI parameters and NOS. We compared graph-theoretic network metrics across weighting schemes and across spatial scales. We then explored structure-function correlations of each weighting scheme with respect to: i) resting-state fMRI functional connectivity data from the same cohort; and ii) population-average measurements of conduction velocities (Lemaréchal et al., 2022) in cortico-cortical evoked potentials from *in vivo* intracranial stereo-electroencephalography (SEEG).

## 2 Results

### 2.1 Network Characteristics

We generated weighted connectomes for n = 66 subjects (age 22–36 years, 29 males) from the Human Connectome Project (HCP) (Van Essen et al., 2013). Connectomes were constructed from tractography data previously used to generate a probabilistic multi-scale white matter atlas (Alemán-Gómez et al., 2022), with four scales of grey matter parcellation (comprising 82, 128, 230 and 460 regions for scale 1, 2, 3 and 4 respectively). Weighted connectomes (Figure 1A) were generated for Standard Model parameters (f and 1/D_e_ ^⟂^) and DTI parameters (FA and 1/RD), in addition to NOS and normalised NOS (normalised by the volume of the grey matter regions at the end points). Since perpendicular diffusivity is expected to have an inverse relationship with white matter connectivity strength (e.g. decreasing with increasing myelination), the inverse of bundle-average values of D_e_^⟂^ and RD were taken.

**Figure 1.**
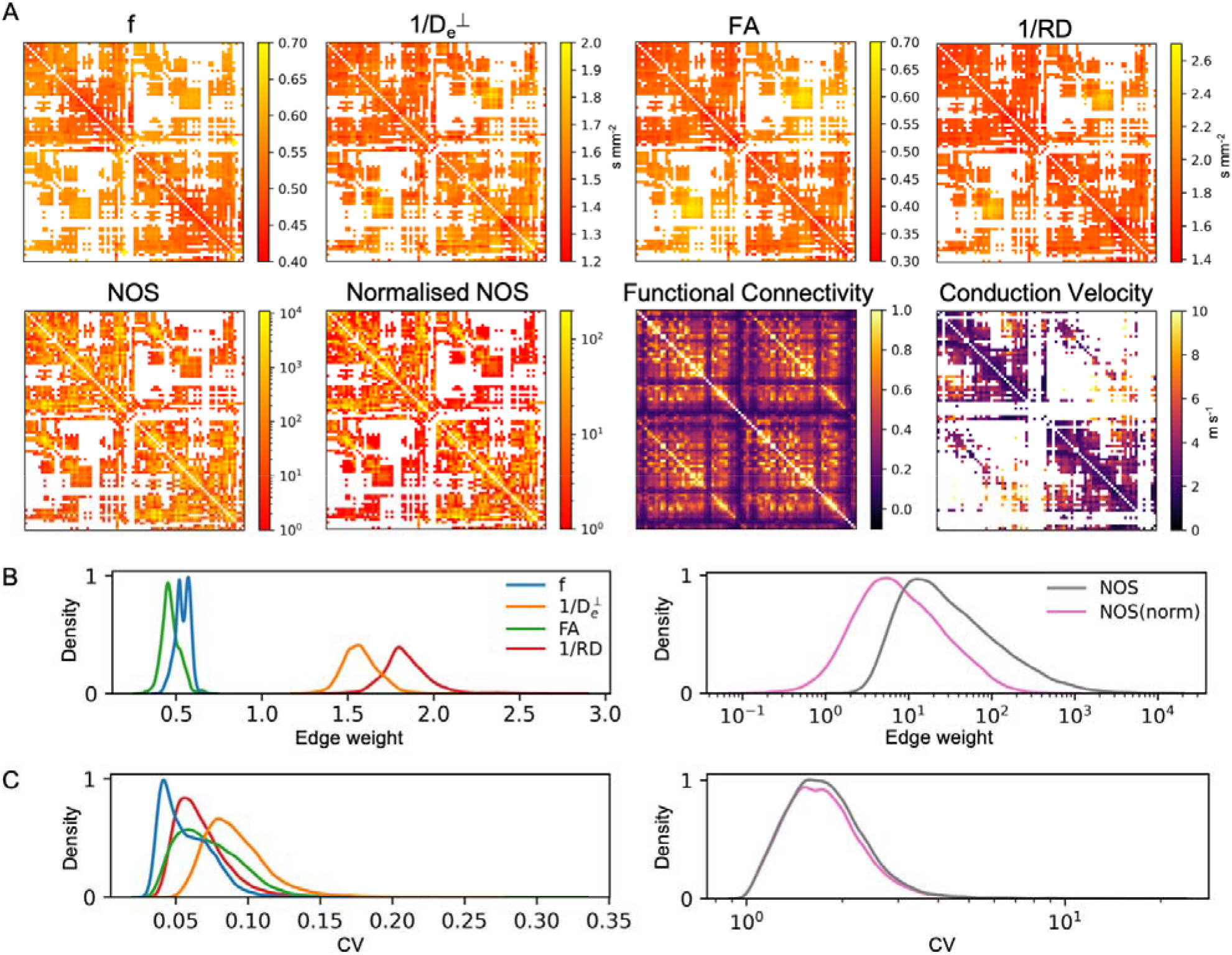
Connectomes for parcellation scale 1 (82 regions). Group-average connectomes are shown for each microstructure weighting (f, 1/D ^⟂^, FA and 1/RD), streamline weighting (NOS and normalised NOS), fMRI functional connectivity, and SEEG conduction velocity (A). Density plots show the distribution of edge weights for structural connectomes (B), and the coefficient of variation (CV) across subjects, calculated for each edge and plotted as a distribution across edges (C).

Each connectome weighting scheme exhibited a distinct distribution of edge weights (Figure 1B). Of the microstructure-weighted connectomes, 1/D_e_ ^⟂^ had the most variation in edge weights across subjects, indicated by a higher coefficient of variation in edge weights across subjects (Figure 1C). There was moderate to high correlation between edge weights in microstructure-weighted connectomes, with the exception of f and 1/D_e_ ^⟂^ which showed minimal correlation at all four parcellation scales (Supplementary Figure 1). There was a strong correlation between NOS-weighted and normalised NOS-weighted connectomes (>0.94), and between FA-weighted and 1/RD-weighted connectomes (>0.85) (Supplementary Figure 1). All connectome weights showed a moderate correlation with bundle length, apart from 1/D_e_ ^⟂^ (Supplementary Figure 1).

The microstructure connectome data derived in this study are publicly available in an updated version of the multi-scale white matter atlas (https://doi.org/10.5281/zenodo.4919131) (Alemán-Gómez et al., 2021).

### 2.2 Small-world Network Properties

For every weighted connectome derived for each subject, we measured the characteristic path length, global efficiency, local efficiency, clustering coefficient, and small-world propensity. Small-world propensity is a normalised measure which is comparable across weighting schemes (Muldoon et al., 2016). The other network metrics were normalised to null networks using two approaches.

First, network metrics were normalised to null networks with randomised edges and weights while preserving strength distributions. Here, the null hypothesis is that the network metric is the same as that of any other network with the same density and strength distribution. This conventional approach to normalising network metrics allows us to verify that connectomes exhibited the trends expected of small-world brain networks. Here, clustering coefficient and local efficiency were higher than random for all weighting schemes and increased with parcellation scale (Figure 2), as expected (Bassett et al., 2011; Zalesky et al., 2010). Global efficiency was not significantly higher than random for any weighting scheme and decreased with increasing parcellation scale. Characteristic path length increased with parcellation scale for all weighting schemes. Despite being low for NOS-weighted connectomes, characteristic path length was not significantly lower than random networks. All weighting schemes exhibited high small-worldness (small world propensity >0.7) across all parcellation scales. For several metrics, the trends for NOS-weighted connectomes deviated from those of microstructure-weighted networks, as expected due to the substantially different edge distribution (Figure 1B).

**Figure 2.**
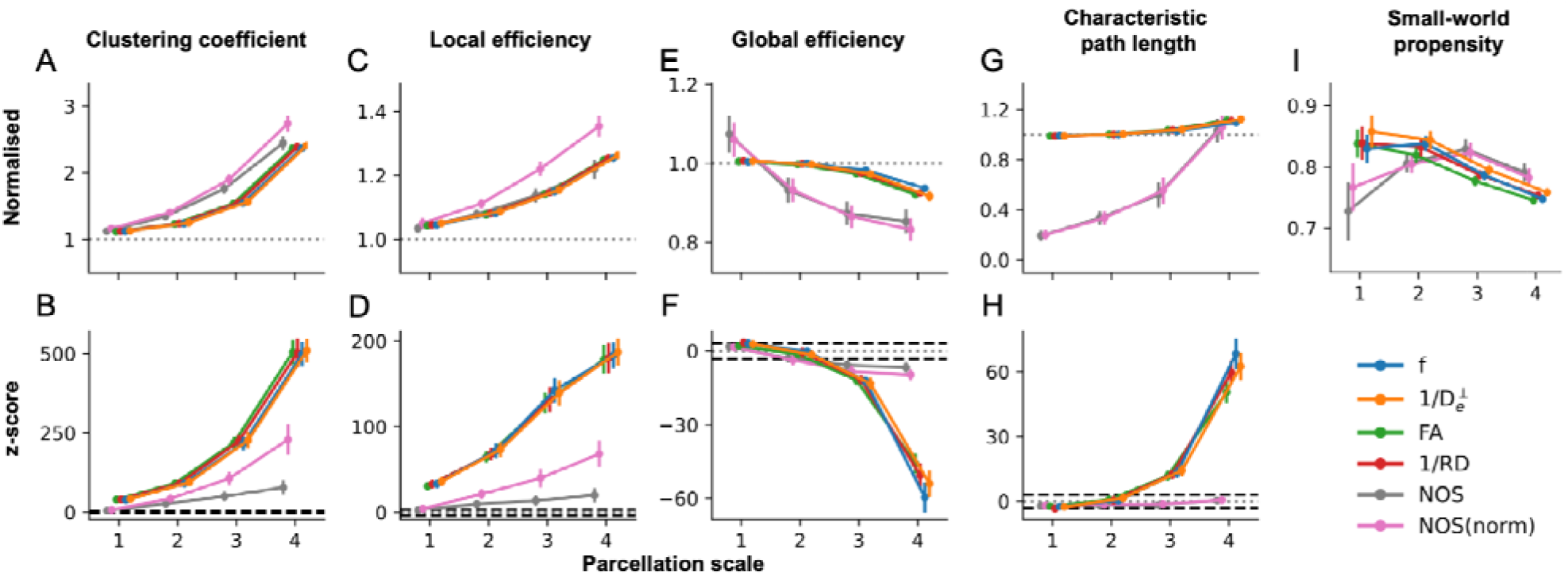
Network metrics across subjects. Normalised metrics and associated z-scores relative to null networks with randomised edges and weights are shown for clustering coefficient (A & B), local efficiency (C & D), global efficiency (E & F) and characteristic path length (G & H). Small-worldness is characterised by small-world propensity (I). Error bars represent the standard deviation across subjects. The mean of the null distribution is indicated by a dotted line on network metric and z-score plots. Significant deviations from the mean of the null distribution (Bonferroni corrected p < 0.05, z > 3.08) are indicated by a dashed line on the z-score plots.

Next, network metrics were normalised to null networks with randomised weights only, while preserving the binary structure. Here, the null hypothesis is that the network metric is the same as that of any other network with the same binary structure and strength distribution. Since the binary structure for a given subject is the same across all weighting schemes, this provides a conservative null distribution revealing the extent to which the network metric is dictated by the specific edge weights sampled from the data, rather than being determined by the binary structure and strength distribution. Thus, this approach allows more direct comparison of the impact of the connectome weighting scheme on the network metrics. Here, connectomes weighted by 1/D_e_ ^⟂^ and normalised NOS gave significantly higher than random clustering coefficient and local efficiency across all scales (Figure 3). Trends in global efficiency and characteristic path length largely reflected those shown above. However, here the global efficiency was significantly higher than random (and characteristic path length was significantly lower) for microstructure-weighted connectomes at scale 1.

**Figure 3.**
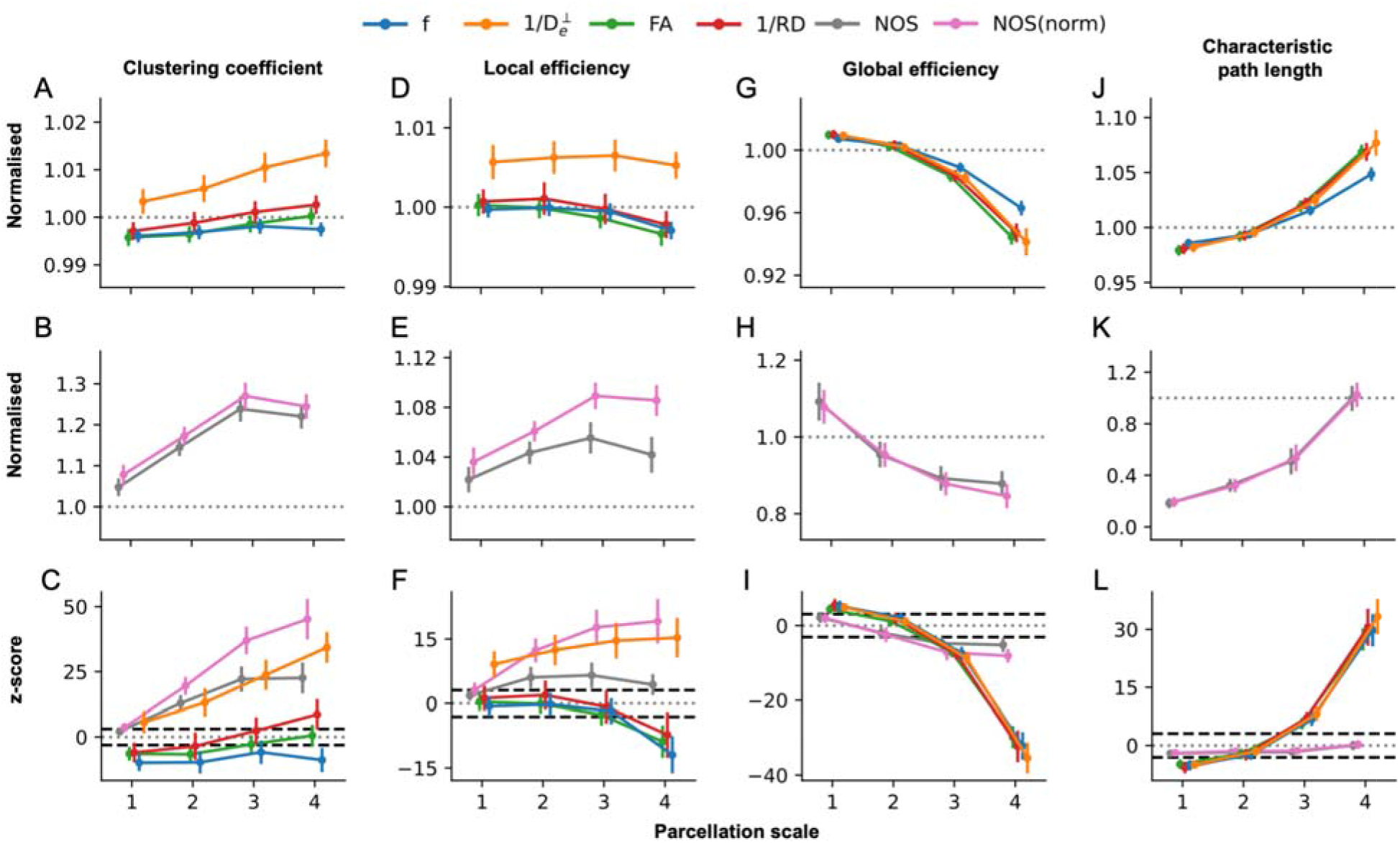
Network metrics relative to null networks with preserved binary structure. Normalised metrics and associated z-scores are shown for clustering coefficient (A-C), local efficiency (D-F), global efficiency (G-I) and characteristic path length (J-L). Metrics for the NOS connectomes are plotted separately for better visualisation of microstructure-weighted connectome metrics. Error bars represent the standard deviation across subjects. The mean of the null distribution is indicated by a dotted line. Significant deviations from the mean of the null distribution (Bonferroni corrected p < 0.05, z > 3.08) are indicated by a dashed line on the z-score plots.

D_e_ ^⟂^ and RD both aim to characterise diffusion perpendicular to the main orientation of white matter fibres, with the key distinction that the diffusion tensor is strongly influenced by fibre dispersion whereas the standard model accounts for multiple fibre populations by fitting a fibre orientation distribution. To investigate the influence of fibre dispersion on network connectivity properties, we measured the local efficiency and clustering coefficient at the node level for group-average 1/D_e_ ^⟂^-weighted and 1/RD-weighted connectomes, and compared these to the mean FA of connections to each node (Supplementary Figure 2). Nodes with lower mean FA (and therefore the highest proportion connections passing through regions of high dispersion) were found to have larger differences in local efficiency (Pearson’s r in the range -0.25 to -0.52, p < 0.001) and clustering coefficient (Pearson’s r in the range -0.31 to -0.52, p < 0.001) between 1/D_e_ ^⟂^-weighted and 1/RD-weighted connectomes (Supplementary Figure 2).

### 2.3 Hub Structure

We then evaluated the hub scores and rich club characteristics of each group-average connectome. Hubs represent densely connected brain regions with a central role in information integration (van den Heuvel & Sporns, 2013; Hagmann et al., 2008). Dense connectivity between hub regions forms a “rich club”, representing the structural core of the connectome (van den Heuvel & Sporns, 2011; Hagmann et al., 2008). Disrupted connectivity to hub nodes is linked to neuropathology, thus it is crucial to be able to identify hub regions and characterise their connectivity (Zalesky & Di Base, 2026; Crossley et al., 2014).

At scale 1, nodes with high hub scores were broadly similar across microstructure-weighted connectomes (Figure 4A), and comprised the precuneus, superior parietal cortex, putamen and thalamus, as expected (van den Heuvel & Sporns, 2011; Hagmann et al., 2008). At finer parcellation scales, high hub scores were also found in regions of the cingulate cortex (Supplementary Figure 3A). At all parcellation scales, f-weighted connectomes had higher hub scores in frontal regions, and microstructure-weighted connectomes tended to have higher hub scores than NOS and normalised NOS-weighted connectomes. These patterns reflected in the Euclidean distance between the hub scores of each connectome (Figure 4B and Supplementary Figure 3B), which show lower distances between connectomes weighted by 1/D_e_ ^⟂^, FA and 1/RD, but slightly higher distances with f-weighted connectomes, and yet higher distances with NOS or normalised NOS-weighted connectomes.

**Figure 4.**
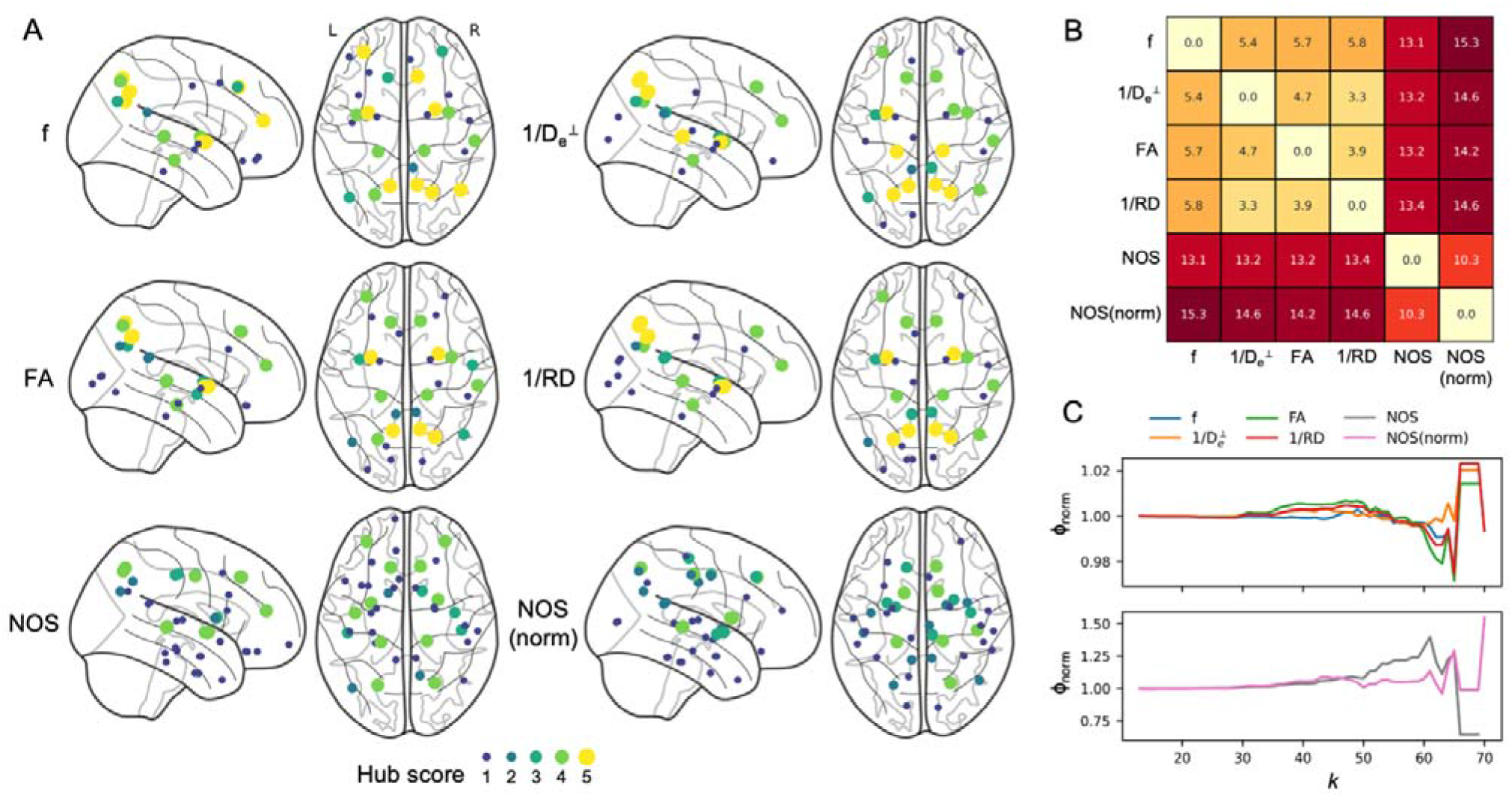
Hub scores and normalised rich club coefficients for parcellation scale 1. Nodes with nonzero hub score are shown for each parameter (A), with the hub score indicated by the colour scale. The pairwise Euclidean distance between hub scores for each parameter is shown in a heatmap (B). Normalised rich club coefficient (□ _norm_) curves are shown for each parameter (C).

The similarities in hub structure between microstructure-weighted connectomes is further reflected in the rich club curves. Connectomes weighted by 1/D_e_ ^⟂^, FA and 1/RD had a rich club regime (□ _norm_> 1) for thresholds in the range 35 < k < 55 at scale 1 (Figure 4C). Similarly, at finer parcellation scales these three connectome weighting schemes exhibited comparable rich club characteristics (Supplementary Figure 4). At scales 1, 2 and 4, 1/D_e_ ^⟂^-weighted connectomes showed a secondary rich club regime at a higher range of thresholds, peaking at k = 64, 82 and 145 respectively. Connectomes weighted by f had a smaller rich club regime at scale 1 (for thresholds in the range 45 < k < 55) and did not have a rich club regime at finer parcellation scales. Compared to microstructure-weighted connectomes, the rich club coefficient for NOS and normalised NOS-weighted connectomes was generally higher and spanned a larger range of thresholds at all scales (Supplementary Figure 4).

### 2.4 Structure-function coupling

To evaluate the functional relevance of each connectome weighting scheme, we measured the structure-function coupling (SFC) of each group-average structural connectome with i) resting-state fMRI functional connectivity data derived from the same population; and ii) SEEG conduction velocity measurements from the F-tract atlas (Lemaréchal et al., 2022; Trebaul et al., 2018). Whole-brain SFC was measured as the Spearman correlation coefficient between structural and functional edge weights, with bundle length included as a regressor, across all edges in the network (Figure 5A, left). For fMRI functional connectivity, regional SFC was measured for each node, using only edges connected to that node (Figure 5A, right). Node-level SFC was not measured for SEEG conduction velocities due to the sparsity of the data limiting the statistical power.

**Figure 5.**
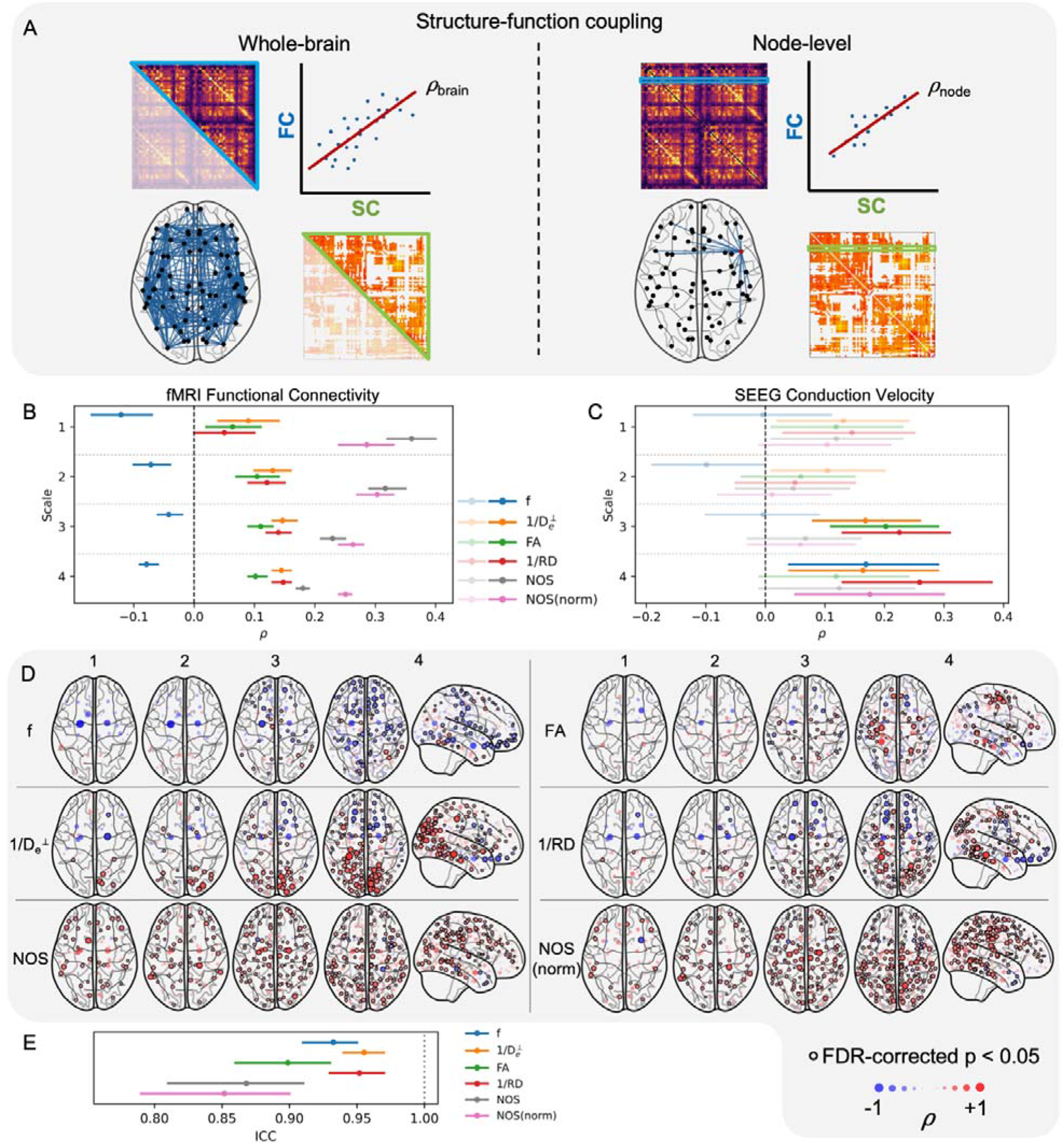
Structure-function coupling. For each weighting scheme at each parcellation scale, the SFC was measured as the Spearman correlation coefficient between structural and functional edge weights, with bundle length included as a covariate. This was measured at the whole-brain level across all edges in the network, and at the node level across edges connected to a given node (A). Whole-brain SFC, with bars extending to the 95% confidence intervals, is shown for fMRI functional connectivity (B) and SEEG conduction velocity (C). Non-significant results (after FDR correction) are shown with reduced opacity. Node-level SFC maps for each parcellation scale show the magnitude and significance of the Spearman correlation for each node, for each connectome weighting scheme (D). Nodes with stronger SFC are represented by larger, more opaque dots, with significance indicated by a black border. Consistency in node-level SFC across parcellation scales is indicated by the ICC with 95% confidence intervals (E).

Whole-brain SFC with fMRI functional connectivity was moderate and significant for NOS-weighted and normalised NOS-weighted connectomes (Figure 5B, Supplementary Figure 5). Coupling decreased with the parcellation scale for NOS-weighted connectomes but was more consistent across scales with normalised NOS-weighted connectomes. For microstructure-weighted connectomes, whole-brain coupling with fMRI functional connectivity was weak but significant at all parcellation scales. Since BOLD fMRI suffers from lower contrast-to-noise ratio in subcortical regions (Kundu et al., 2012), we also measured whole-brain SFC with only cortico-cortical connections (whole-cortex SFC). For f-weighted connectomes, the whole-cortex SFC was not significant at scales 1 and 2, and was weakly positive for scale 3 (Supplementary Figure 6). This indicates that the negative SFC when considering all brain regions was driven by weak functional connections to the subcortical structures, which are in proximity to areas of high axon density. At scale 4, whole-cortex SFC remained weakly negative, possibly due to remaining deep brain regions with abnormally low functional connectivity, such as the entorhinal cortex and parahippocampal gyrus, where the low contrast-to-noise ratio is compounded by the small volume of regions at this parcellation scale. For all other connectome weighting schemes, the whole-cortex SFC (Supplementary Figure 6) was comparable to or higher than the whole-brain SFC (Figure 5B), indicating these relationships are robustly exhibited across the whole brain.

Whole-brain SFC with SEEG conduction velocity was weak for all parameters but significant for 1/D_e_ ^⟂^, FA and 1/RD at scale 3, and for f, 1/D_e_ ^⟂^, 1/RD and normalised NOS at scale 4 (Figure 5C, Supplementary Figure 7). Before FDR correction, there was significant coupling with 1/D_e_ ^⟂^, FA, 1/RD and NOS at scale 1, and with 1/D_e_ ^⟂^ at scale 2, but these did not survive multiple comparisons correction due to the high uncertainty resulting from the sparsity of the SEEG data.

Node-level SFC with fMRI functional connectivity increased with parcellation scale for all parameters (Figure 5D). Of the microstructure-weighted connectomes, 1/D_e_ ^⟂^ had the most widespread significant coupling. NOS-weighted and normalised NOS-weighted connectomes had high coupling across widespread brain regions. To evaluate the similarity in regional SFC patterns across parcellation scales, the node-level SFC values at scales 2, 3 and 4 were z-transformed and averaged across regions belonging to the same scale 1 region. The intraclass correlation coefficient (ICC) was then measured across z-transformed coupling values to give a measure of similarity across scales for each parameter. The consistency across spatial scales was highest for connectomes weighted by 1/D_e_ ^⟂^ and 1/RD (Figure 5E).

## 3 Discussion

Structural connectomes typically aim to characterise the strength of white matter connections between brain regions in order to capture topological organisation and potential for transmission of functional information across the brain. In this work, we investigated the use of microstructure parameters derived from the Standard Model of diffusion in white matter as connectome weights, with the goal of leveraging their enhanced specificity to the microstructural properties of the underlying tissue compared to other microstructure models or signal representations (Coronado-Leija et al., 2024; Jelescu et al., 2016b; Liao et al., 2024), ultimately enabling integration of a wider range of complementary white matter connectivity properties into brain network analysis. We evaluated connectomes at multiple spatial scales in terms of their ability to capture expected global properties of brain network organisation, and their relationship with functional measures from multiple modalities, in comparison with weights derived from DTI and NOS.

First, we validated the ability of each weighting scheme to capture expected properties of small-world brain networks (Bassett & Bullmore, 2017; Bassett & Sporns, 2017; Bullmore & Bassett, 2011; Kaiser, 2011). All weighting schemes captured significant clustering and local efficiency, indicating a high level of network segregation. While microstructure-weighted networks had high path length and low global efficiency at finer parcellation scales, this was sufficiently balanced by high segregation to result in a high small-worldness for all weighting schemes. All weighting schemes also reproduced previously reported trends of increasing clustering coefficient, path length and local efficiency, and decreasing global efficiency, with increasing number of nodes (Bassett et al., 2011; Zalesky et al., 2010).

Since the edge weight distribution influences network metrics, appropriate normalisation to null network models is essential for comparison of network metrics between connectomes with different weighting schemes (Váša & Mišić, 2022). Normalisation to null networks with randomised binary structure and edge weights revealed topological properties which are not explained by the strength distribution alone, consistent with previous works (Bassett & Bullmore, 2017; Bullmore & Bassett, 2011). However, many network properties were similar across weighting schemes, particularly for microstructure-weighted connectomes, suggesting the binary structure has a substantial influence on the network metrics. Since the binary structure of the network is the same across different weighting schemes for a given subject, we sought to further evaluate the influence of each weighting scheme on network properties using null network models with randomised edge weights while preserving binary structure. In this case, only connectomes weighted by 1/D_e_ ^⟂^ and normalised NOS had significantly higher clustering coefficient and local efficiency than the null distribution at all parcellation scales. This shows that the high segregation in networks with these weighting schemes does not arise solely due to the binary structure and strength distribution, but also depends on the specific arrangement of edge weights (Liégeois et al., 2021). For all other weighting schemes, comparable network metrics were found in randomised networks with the same binary structure and strength distribution. Thus, connectomes weighted by 1/D_e_ ^⟂^ and normalised NOS capture features of non-random brain network organisation which are not captured by the other weighting schemes.

A likely explanation for the enhanced sensitivity of 1/D_e_ ^⟂^-weighted connectomes to local organisation is the improved specificity to diffusion perpendicular to fibre bundles which is gained by fitting the fibre orientation distribution, contrary to RD which remains susceptible to confounding effects of fibre dispersion (Coelho et al., 2022; Liao et al., 2024). Supporting this, we showed that the nodes with the lowest mean FA, representing the nodes with the highest proportion of connections passing through regions of high fibre dispersion, had the biggest differences between 1/D_e_ ^⟂^ and 1/RD in the local connectivity properties.

Beyond small-world metrics of integration and segregation, we evaluated the hub structure and rich club properties of microstructure-weighted connectomes. Each weighting scheme captured a broadly consistent hub structure and comparable rich club characteristics, indicating that sensitivity to these topological properties is preserved across microstructure parameters. One exception is the secondary rich club regime observed in 1/D_e_ ^⟂^-weighted connectomes at some scales, possibly reflecting increased sensitivity to the organisation of the most densely interconnected hub subnetworks.

Together with the analysis of small-world network properties, these findings suggest that large-scale topological network properties are preserved across microstructure-weighted connectomes, while sensitivity to local network segregation maybe enhanced with 1/D_e_ ^⟂^-weighted connectomes. Further study is needed to determine whether these local features captured by 1/D_e_ ^⟂^-weighted connectomes provide improved sensitivity to intersubject differences (e.g. in predicting age or cognition) and to perturbations in network connectivity due to disease or disorder (Liu et al., 2017).

Next, we evaluated the functional relevance of each connectome weighting scheme with respect to brain function measured with different modalities at different temporal scales. Resting-state BOLD fMRI functional connectivity measures synchronous patterns of activity on a timescale of several seconds (Biswal et al., 1995; Horwitz, 2003), while SEEG conduction velocity provides a measure of the propagation of electrical signals on a timescale of milliseconds (Lemaréchal et al., 2022). As conduction velocities were derived from delay measurements below 50 ms, they are expected to reflect predominantly direct white matter connections (though may also be influenced by synaptic transmission delays and indirect connections) (Avalos-Alias et al., 2026; Lemaréchal et al., 2022; Trebaul et al., 2018). At the whole-brain level, we reproduced the widely reported correlation between NOS-weighted connectomes and fMRI functional connectivity (Fotiadis et al., 2024; Honey et al., 2009; Suárez et al., 2020). However, there was lower whole-brain coupling of NOS-weighted connectomes with SEEG conduction delays. NOS lacks specificity to any given microstructural properties, and is biased by tractography parameters in addition to features of the fibre path such as length, curvature and degree of branching (Jones, 2010; Jones et al., 2013). While NOS captured connectivity characteristics reflective of the long-timescale dynamics of fMRI functional connectivity, it appears to lack sensitivity to the microstructural features which relate to axonal conduction velocities (Asadi et al., 2025; Caminiti et al., 2013; Drakesmith et al., 2019; Mancini et al., 2021). Microstructure-weighted connectomes had relatively low whole-brain SFC but exhibited similar trends across parcellation scales with both fMRI functional connectivity and SEEG conduction delays. Notably, only 1/D_e_ ^⟂^-weighted connectomes had significant coupling at all scales and modalities before multiple comparisons correction. Evidence from demyelinating disorders suggests D_e_ ^⟂^ is sensitive to myelination and g-ratio (Jelescu et al., 2016b; Falangola et al., 2014; Fieremans et al., 2012). Thus, this trend aligns with previous studies relating myelin-sensitive measures to conduction velocities (Drakesmith et al., 2019; Mancini et al., 2021) and functional connectivity (Nelson et al., 2026). Counterintuitively, f-weighted connectomes had negative SFC with fMRI functional connectivity. By comparison with SFC across cortico-cortical connections, which did not exhibit a consistent trend across all parcellation scales, we determined that this pattern was driven by a subset of weak functional connections in deeper brain areas with lower contrast-to-noise ratio. Notably, SFC with all other weighting schemes was not dependent on the inclusion of subcortical regions.

There is increasing interest in regional differences in SFC (Fotiadis et al., 2024; Gu et al., 2021; Suárez et al., 2020; Vázquez-Rodríguez et al., 2019). Therefore, we explored node-level SFC with fMRI functional connectivity. All weighting schemes showed more widespread coupling at finer parcellation scales, as expected due to the larger number of connections inflating the statistical power. The node-level SFC patterns were qualitatively similar between 1/D_e_ ^⟂^ and 1/RD, with both resembling previously reported unimodal-transmodal gradients characterised by high SFC in primary visual and motor cortices (Vázquez-Rodríguez et al., 2019). 1/D_e_ ^⟂^ showed more widespread association, possibly reflecting the enhanced specificity of D_e_ ^⟂^ which factors out orientation dispersion (Coelho et al., 2022; Liao et al., 2024). Connectomes weighted by 1/D_e_ ^⟂^ and 1/RD had the highest consistency across parcellation scales. In the case of whole-brain SFC with fMRI functional connectivity, the scale dependence of NOS-weighted connectomes (indicated by diminishing SFC with spatial scale) was alleviated in normalised NOS-weighted connectomes. However, the scale-dependence of node-level SFC was not improved by normalising the NOS-weighted connectomes. This scale-dependence highlights the arbitrary nature of NOS as a measure of structural connectivity. Although normalisation by the node volume can alleviate the bias associated with streamline seeding, there are several other biases (described above) which may have a different influence at different scales. Conversely, connectome weights derived from tractometry provide objective parameter values which can be related to microstructural properties of the underlying tissue (Coronado-Leija et al., 2024; Jelescu et al., 2016b; Liao et al., 2024).

This work builds upon previous studies employing microstructure-sensitive parameters for connectomics (Boshkovski et al., 2021; Cavallo et al., 2026; Gast & Assaf, 2024; Mancini et al., 2018; Nelson et al., 2023). Connectomes weighted by myelin-sensitive measures, including g-ratio and R1, have been shown to provide complementary information to NOS-weighted connectomes (Boshkovski et al., 2021; Mancini et al., 2018). However, lacking detailed comparisons with other tractometry-derived connectome weights, it is unclear whether such properties were similarly captured by other microstructure metrics. A more extensive evaluation of connectome weighting schemes (Nelson et al., 2023) revealed distinct patterns among connectomes weighted by R1, intracellular volume fraction (ICVF) from NODDI, DTI metrics (FA and RD), and NOS, in addition to weighted streamline measures from SIFT2 (Smith et al., 2015) and COMMIT (Daducci et al., 2014). Similar to our findings, they identified a divergence between streamline-based connectome weights and those derived from tractometry, but with both exhibiting small-world characteristics. For ICVF, FA and RD, they reported no correlations with functional connectivity after regressing length, whereas we found significant (albeit weak) correlations with 1/D_e_ ^⟂^, FA and 1/RD, possibly due to the large amount of fMRI data providing more robust estimates of functional connectivity in our study (4 sessions per subject, >14 minutes per session, as provided by the HCP).

Notably, our findings from f-weighted connectomes differed from other metrics in terms of the negative whole-brain SFC (attributed to a lack of robust cortical SFC) and a lack of rich club regime at all scales. Previous results in connectomes weighted by ICVF (a conceptually equivalent metric from the NODDI model) similarly reported no correlation with functional connectivity when regressing length, but did identify a rich club regime at multiple parcellation scales (Nelson et al., 2023). These differing rich club results between f and ICVF highlight the differences between the NODDI model and the Standard Model; crucially, NODDI relies on a tortuosity model which may introduce biases in the presence of myelin or in areas of tight packing (Jelescu et al., 2015; Fieremans et al., 2008) and imposes a one-to-one relationship between D_e_ ^⟂^ and ICVF. This inability to distinguish trends in ICVF from D_e_ ^⟂^ represents a key distinction between connectomes weighted by f and those weighted by ICVF, which may capture some confounding effects of f and D_e_ ^⟂^. Connectome weights which use estimates of g-ratio derived from NODDI parameters (in conjunction with magnetisation transfer data) may also inherit these limitations, as previously acknowledged (Mancini et al., 2018).

In this work, the sensitivity of 1/D_e_ ^⟂^-weighted connectomes to local organisation may be due to a combination of resolving multiple fibre populations, sensitivity to myelination, and sensitivity to fibre density, but cannot be attributed to any single one of these factors. Due to its short T2, myelin (or myelin water between the wrappings of the sheath), it is effectively “MRI-invisible” at echo times used in typical clinical diffusion protocols. As such, D_e_ ^⟂^ should be considered a proxy for myelination rather than a direct measure. Furthermore, evidence for the sensitivity and specificity of D_e_ ^⟂^ to myelination was demonstrated in the context of de- and re-myelination within the corpus callosum (Jelescu et al., 2016b; Fieremans et al., 2012; Falangola et al., 2014), therefore it remains unclear whether this specificity extends to differences in myelination between regions of healthy tissue across the brain. These considerations highlight the need to nonetheless move towards multimodal acquisition strategies that combine diffusion MRI with complementary myelin-sensitive measures such as R1. Although these measures are not yet widely available in large open databases, and suffer from their own limitations to specificity (Hutchinson et al., 2024; Harkins et al., 2016; Stüber et al., 2014), their integration with diffusion-based measures may further augment this picture and provide a more comprehensive view of the microstructural basis of macroscale brain organisation.

In addition to evaluating the utility of Standard Model parameters as connectome weights, we provide these weighted connectomes as a supplement to the publicly available multi-scale white matter atlas (Alemán-Gómez et al., 2022). This resource can serve as a reference for the connectome in a healthy population for brain network analysis studies, or as a structural substrate for biophysical modelling of brain network dynamics (Breakspear, 2017; Cabral et al., 2017; Honey et al., 2007, 2009; Seguin et al., 2023a). These modelling approaches may further bridge the gap between structure and function and determine whether sensitivity of 1/D_e_ ^⟂^-weighted connectomes to local organisation can be leveraged to enhance biophysical models of whole-brain network dynamics.

### Limitations

As described above, the inability of diffusion MRI to directly measure myelination impedes our capacity to attribute these characteristics of macroscale brain organisation to myelin specifically, which requires further work integrating multimodal acquisitions. Additionally, evidence for the sensitivity and specificity of Standard Model parameters to underlying microstructural features relies on specific regions of interest (as is inherent to validation studies using electron microscopy or histology), thus may not generalise well to brain-wide white matter with more complex geometry (Jelescu et al., 2016b; Fieremans et al., 2012). However, there is increasing evidence that the Standard Model offers reliable parameter estimates even in regions of high fibre dispersion (Coronado-Leija et al., 2024; Nguyen-Duc et al., 2026).

Another key limitation of this work is that the SEEG conduction delay measurements were acquired from patients with temporal lobe epilepsy. Thus, these may not accurately reflect the conduction velocities in the healthy population from which the rest of the data were derived. This, along with the sparsity of these data, likely contributed to the high uncertainty in the SFC values with SEEG data. However, the dataset was specifically curated to exclude pathological zones and has been demonstrated to generalise to healthy connectivity patterns (Lemaréchal et al., 2022).

Finally, we measured functional connectivity from fMRI using Pearson correlation coefficients. Instead, using partial correlation to measure functional connectivity has been shown to provide stronger coupling with structural connectivity (Liégeois et al., 2020), thus may further accentuate the differences in SFC between weighting schemes.

## Conclusions

Connectome weights derived from D_e_ ^⟂^ from the Standard Model of diffusion in white matter capture meaningful aspects of brain network organisation beyond those reflected by DTI or tractography-based measures, and correlate with measures of brain function from multiple modalities and at multiple spatial scales. Thus, connectomes weighted by biophysical model parameters could provide higher sensitivity to brain network changes, for example those resulting from disease or disorder. Future studies could offer a more detailed characterisation of regional structure-function coupling patterns across microstructure parameters.

## 4 Methods

### 4.1 Data

DWI and tractography data used in this study were the same as those used to generate the probabilistic multi-scale white matter atlas described in Alemán-Gómez et al., (2022). These data consisted of n = 66 subjects (age 22–36 years, 29 males) from the Human Connectome Project (HCP) (Van Essen et al., 2013). We also obtained resting-state fMRI data for the same subjects for investigation of structure-function coupling (SFC). Each subject was scanned on a Siemens 3 T Skyra scanner in Washington University or University of Minnesota. The original HCP protocols were approved by the Institutional Review Board at Washington University in St. Louis, and all participants provided written informed consent (Van Essen et al., 2013).

#### 4.1.1 Structural MRI

T1-weighted sagittal images were acquired using a Magnetization-Prepared Rapid Acquisition Gradient Echo (MPRAGE) sequence with 3D inversion recovery, echo time (TE) □ = □ 2.14 ms, repetition time (TR) □ = □ 2400 ms, inversion time □ = □ 1000 ms, flip angle = □ 8°, bandwidth □ = □ 210 Hz per pixel, echo spacing (ES) □ = □ 7.6 ms, gradient strength □ = □ 42 mT m^-1^, field of view (FoV) □ = □ 180 × □ 224 × □ 224 mm^3^, and voxel size □ = □ 0.7 × □ 0.7 × □ 0.7 □ mm^3^.

#### 4.1.2 Diffusion MRI

DWI data were acquired with a multi-slice echo planar imaging (EPI) sequence with multi-band (MB) excitation and multiple receivers were acquired with TE □ = □ 89.5 ms, TR □ = □ 5520 ms, flip angle □ = □ 78°, refocusing flip angle □ = □ 160°, bandwidth □ = □ 1488 Hz per pixel, MB factor □ = □ 3, ES □ = □ 0.78 ms, gradient strength □ = □ 100 mT m^-1^, FoV □ = □ 210 × 180 □ × □ 138 mm^3^, voxel size = □ 1.25 □ × □ 1.25 □ × □ 1.25 □ mm^3^ and b-values □ = □ 0, 1000, 2000 and 3000 s mm^-2^. A full dMRI session included six runs, comprising three different gradient tables, each acquired with right-to-left and left-to-right phase encoding. Each gradient table included approximately 90 diffusion weighting directions plus six interspersed b = 0 volumes.

#### 4.1.3 Functional MRI

Resting-state fMRI data were acquired with a gradient-echo EPI sequence with TE □ = □ 33.1 ms, TR □ = □ 720 ms, flip angle □ = □ 52°, bandwidth □ = □ 2290 Hz per pixel, MB factor □ = □ 8, ES □ = □ 0.58 ms, gradient strength □ = □ 100 mT m^-1^, FoV □ = □ 208 □ × □ 180 □ × □ 144 mm^3^, voxel size □ = □ 2.0 □ × □2.0 □ × □ 2.0 mm^3^. Four resting-state runs were acquired over two sessions (one right-to-left phase encoding, one left-to-right phase encoding per session), with 1200 frames (14 minutes 33 seconds) per run.

#### 4.1.4 Conduction Delays

Conduction delay measurements were obtained from the F-tract atlas (Lemaréchal et al., 2022; Trebaul et al., 2018). These were derived from cortico-cortical evoked potentials measured from intracranial SEEG recordings in patients with pharmaco-resistant focal epilepsy, while awake and at rest during resective surgery. Data were aggregated from 21 centres, with a total of 1307 implantations in 1236 patients: 602 male (mean age (s.d.) 25 (14) years); 628 female (mean age (s.d.) 26 (13) years); 6 unknown (mean age (s.d.) 30 (10) years). The precise signal processing methodology is described in earlier work (Seguin et al., 2023b) and outlined briefly below. First, a machine learning algorithm (supervised by trained personnel) was used to identify bad channels. According to standard procedure, signals were referenced to the bipolar montage to minimise volume conduction effects. Following removal of short stimulation artefacts (<4 ms) by local signal blanking and interpolation, band-pass filtering was applied with cutoff frequencies of [1, 45] Hz. All stimulation pulses within one run were then averaged, baseline-corrected, and normalised to [-200, -10] ms before stimulation. Averaged responses were considered significant if their absolute value exceeded a z-score of 5 within a [0, 50] ms time window. This 50 ms cutoff was chosen to increase specificity to direct connections, avoiding contributions of axonal propagation through multiple regions. Responses were obtained for 10,551,456 recordings from 90,653 stimulations with mean (s.d.) current intensity 3.4 (1.3) mA, mean (s.d.) pulse width 1 (0.5) ms, and frequency 1 Hz. Stimulations were biphasic in 75%, and monophasic in 25% of instances.

To characterise conduction delay, we selected the peak delay, defined as the time between stimulation and the peak of the first significant component of the response. Population-average measurements of peak delays were mapped onto each parcellation scale. Only connections with at least 50 measurements from at least 3 patients were considered. Since the delay measurements are asymmetrical, the bidirectional average was taken. Conduction velocity was calculated by dividing tract length by conduction delay, using group-average tract length measurements from the multi-scale white matter atlas (Alemán-Gómez et al., 2022).

### 4.2 Structural Connectome Construction

DWI preprocessing steps applied as part of the HCP processing pipeline were as follows: normalisation of the b0 image intensity across runs; removal of EPI distortions, eddy-current-induced distortions, and subject motion; gradient-nonlinearity correction; registration to the T1-weighted structural image resampled to 1.25 mm resolution. For more details on these preprocessing steps, see Glasser et al., (2013). The process for generating weighted connectomes for each subject is described below and summarised in Figure 6.

**Figure 6.**
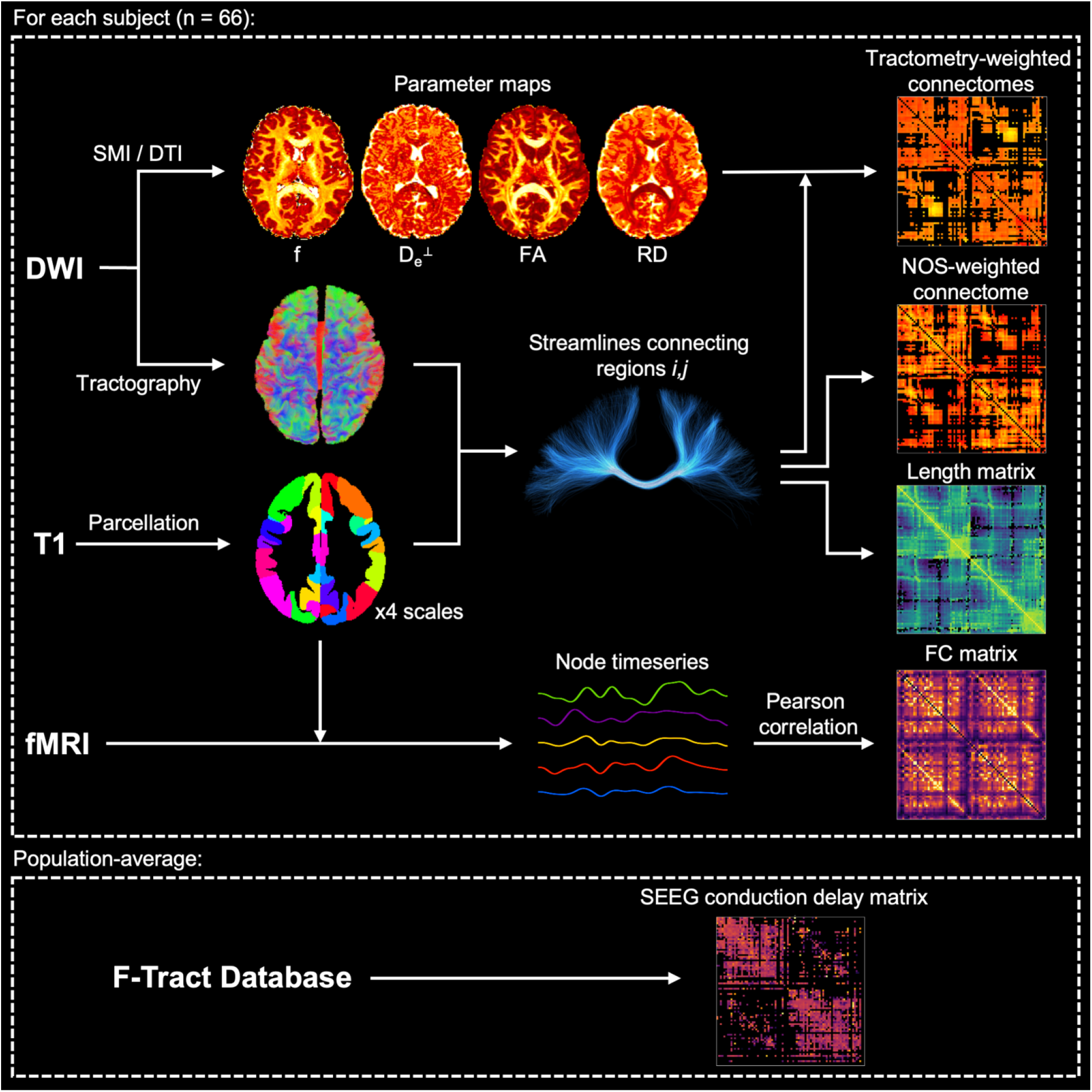
Connectome construction. For each subject, connectomes weighted by diffusion parameters were generated for each parcellation scale by measuring the average of a given parameter along each streamline and averaging across streamlines in a given bundle. Number of streamlines and bundle length were also measured for each subject. Functional connectomes were generated by measuring the Fisher z-transformed Pearson correlation between the average BOLD signal in each pair of regions. A population-average matrix of SEEG conduction delays was obtained from the F-tract database (Lemaréchal et al., 2022).

#### 4.2.1 Connectivity

Connectomes were derived from tractography data previously used to generate a probabilistic multi-scale white matter atlas (Alemán-Gómez et al., 2022). Processing steps are described briefly below, with further details provided in Alemán-Gómez et al., (2022).

The T1-weighted image for each subject was segmented into four scales of grey matter parcellation (Cammoun et al., 2012; Fischl et al., 2002). At scale 1, the cortical parcellation corresponds to the Desikan-Killiany atlas provided in the HCP database (Desikan et al., 2006; Glasser et al., 2013). Scales 2, 3 and 4 provide progressively finer subdivisions of the scale 1 atlas. Cortical parcellations were combined with a subcortical segmentation comprising the striatal structures (caudate nucleus, putamen and nucleus accumbens), globus pallidum, amygdala, hippocampus, brainstem, and the thalamus subdivided into seven nuclei (Battistella et al., 2017). For the present study, only grey matter regions were considered (i.e. the brainstem was removed), and thalamic nuclei were combined to give a single thalamus region. This resulted in 82, 128, 230 and 460 regions for scale 1, 2, 3 and 4 respectively.

White matter bundles between pairs of grey matter regions were estimated from DWI using constrained spherical deconvolution (Descoteaux et al., 2009; Tournier et al., 2007) and anatomically-constrained particle filtering tractography (Girard et al., 2014). Tractography was seeded from the interface between grey matter and white matter with 30 seeds per voxel, and only streamlines with length between 20 mm and 200 mm were kept. For each parcellation scale, streamlines were assigned to bundles connecting pairs of regions and filtered to remove outliers (streamlines taking anatomically implausible paths) (Côté et al., 2015; Cousineau et al., 2017). To suppress spurious connections and ensure consistency across subjects, bundles present in less than 80% of subjects were removed.

#### 4.2.2 Edge Weights

Standard Model parameters were fitted to DWI data using the Standard Model Imaging (SMI) toolbox (Coelho et al., 2022). Fitting the Standard Model required a map of noise standard deviation (σ) characterising the DWI data. Therefore, we used the *dwidenoise* tool from MRtrix (Tournier et al., 2019) to estimate the σ map from the raw DWI data for each subject, which was then transformed to the same space as the preprocessed DWI data. To avoid degeneracy due to over-parameterising the model, the free-water compartment was neglected (i.e. setting f_w_ = 0) based on evidence that this improves the sensitivity and specificity of the remaining standard model parameters in a standard multi-shell sequence (Coelho et al., 2024). Diffusion tensor imaging (DTI) parameters were fitted to DWI data using TMI in Designer-v2 (Chen et al., 2024).

For connectome edge weights, we used Standard Model parameters f and D_e_ ^⟂^, in addition to DTI parameters FA and RD. For each parameter, the average along each streamline was measured, then the edge weight was defined as the mean value across all streamlines in the bundle. For D_e_ ^⟂^ and RD, the inverse of bundle-average values were taken. In addition to these tractometry-derived edge weights, NOS-weighted connectomes were generated with edge weights defined as the number of streamlines belonging to each fibre bundle. Normalised NOS-weighted connectomes were also generated by dividing each bundle count by the total volume of the corresponding pair of grey matter regions.

### 4.3 Network Properties

Network metrics were calculated using the brain connectivity toolbox (Rubinov & Sporns, 2010). For each weighting scheme, we measured several network metrics for each subject:

^⍰^Characteristic path length: the average shortest path length between all pairs of nodes in the network, providing a measure of network integration. Note that ‘path lengths’ do not correspond to physical distances, but are defined inversely to edge weights, such that a strong connection gives a short path length.
^⍰^Global efficiency: the average inverse shortest path length between all pairs of nodes. This is a similar measure of network integration to characteristic path length, but in this case the measure is primarily influenced by short paths (strong connections).
^⍰^Local efficiency: measured for a given node as the average inverse shortest path between neighbours of that node, via paths only containing neighbours of that node. This is averaged across nodes to give an average local efficiency for the whole network. This measure of network segregation reflects the efficiency of connectivity in subgroups of interconnected nodes.
^⍰^Clustering coefficient: for a given node this reflects the proportion of a node’s neighbours which are also neighbours of each other. For weighted graphs this is calculated as the average geometric mean of triangles around the node. Averaging across all nodes gives a measure of network segregation reflecting the prevalence of clustered connectivity around nodes.
^⍰^Small-world propensity: small-worldness characterises the degree to which a network exhibits both high integration *and* segregation (Bassett & Bullmore, 2006). A network is considered ‘small-world’ if it has a high clustering compared to random networks, whilst having comparable (or only slightly higher) path length. Small-world propensity is calculated from measures of clustering and path length normalised to lattice networks and random networks, providing a continuous scalar metric of small-worldness which is independent of edge density (Bassett & Bullmore, 2017; Muldoon et al., 2016).

To offer a meaningful comparison between different weighting schemes, the first four of these network metrics were normalised to randomised networks. Null networks were generated with two approaches (Figure 7): i) rewiring edges and randomising edge weights while preserving the strength distribution; and ii) randomising weights only, while preserving the binary structure and strength distribution. For each normalisation approach, 100 null networks were generated per subject, then the network metric was expressed in terms of i) a normalised metric calculated by dividing the original metric by the mean of the null distribution; and ii) a z-score relative to the null distribution.

**Figure 7.**
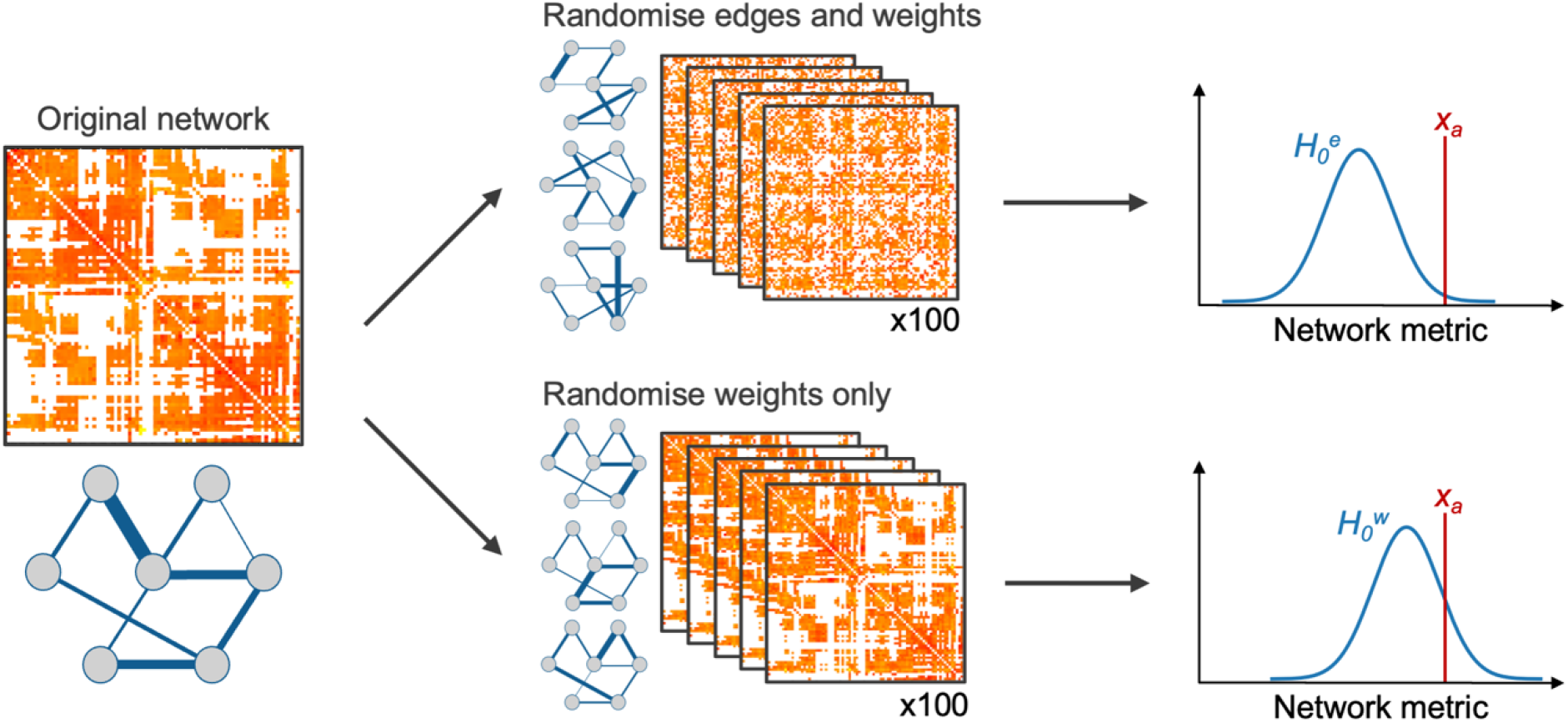
Normalisation of network metrics. Two null network models were used: one randomising the edges and weights (top), and one randomising only weights while preserving the binary structure (bottom).

Hub scores for each node were derived by adding one to nodes in the highest 20% for node strength, betweenness, eigenvector centrality, and closeness, and in the lowest 20% for clustering coefficient (van den Heuvel et al., 2010; van den Heuvel & Sporns 2013). Rich club curves were calculated by computing the rich club coefficient (□) at each degree threshold (k) (van den Heuvel & Sporns, 2011). Rich club coefficients were normalised to 1000 null networks, computed using simulated annealing whilst preserving the degree and strength distribution (Liu et al., 2025; Milisav et al., 2025; Mišić et al., 2015).

### 4.4 Functional Connectomes

Resting-state fMRI functional connectomes were constructed in order to measure SFC with each of the weighted structural connectomes. Resting-state fMRI preprocessing steps applied as part of the HCP processing pipeline were as follows: gradient distortion correction; motion correction to a single-band reference image; fieldmap-based distortion correction; registration to the T1-weighted structural image and resampling to 1.25 mm resolution; highpass filtering (0.01 Hz cutoff); and independent component analysis (ICA)-based artifact removal using FSL FIX, to remove non-neural components, including regression of motion parameters (Griffanti et al., 2014; Salimi-Khorshidi et al., 2014; Smith et al., 2013). At each parcellation scale, the BOLD signal was averaged within each region, then demeaned, low-pass filtered (0.1 Hz cutoff) and concatenated across four runs. A functional connectome was generated for each subject at each parcellation scale by measuring the Fisher z-transformed Pearson correlation between each pair of regions. This was averaged across subjects to obtain a group-average functional connectome for each parcellation scale.

### 4.5 Structure-Function Coupling

We measured the SFC of each group-average structural connectome with resting-state fMRI functional connectivity and with SEEG conduction velocity. Whole-brain SFC was measured as the Spearman correlation coefficient between structural and functional edge weights, with bundle length included as a regressor, across all edges in the network (Figure 5A, left). Regional SFC was measured for each node, using only edges connected to that node (Figure 5A, right).

### 4.6 Statistical Analysis

Network metrics were calculated for each structural connectome at each scale. Using both null network approaches described above, metrics were plotted as normalised values and as z-scores. Significant deviations from the null distribution were defined by a z-score corresponding to Bonferroni-corrected p < 0.05 (z > 3.08).

Whole-brain SFC was measured for each group-average structural connectome at each scale, and plotted as the correlation and 95% confidence interval. This was measured for both fMRI functional connectivity and SEEG conduction velocity. Node-level SFC with fMRI functional connectivity was measured for each region. For both node-level and whole-brain SFC, p-values were measured using permutation testing with 5000 permutations, and FDR-correction applied.

To evaluate the similarity in regional SFC patterns across parcellation scales, the node-level SFC values at scales 2, 3 and 4 were z-transformed and averaged across regions belonging to the same scale 1 region. The intraclass correlation coefficient (ICC) was then measured across z-transformed coupling values to give a measure of similarity across scales for each parameter. Node-level SFC analysis was not performed for SEEG conduction velocity due to the sparsity of the data limiting the statistical power.

## Acknowledgements

The authors would like to thank Santiago Coelho for advice regarding Standard Model parameter estimation. This work was funded by the Swiss National Science Foundation Sinergia grant no. 209470, Eccellenza Fellowship no. 194260 and Swiss Secretariat for Research and Innovation award no. MB22.00032. Data were provided in part by the Human Connectome Project, WU-Minn Consortium (Principal Investigators: David Van Essen and Kamil Ugurbil; 1U54MH091657) funded by the 16 NIH Institutes and Centers that support the NIH Blueprint for Neuroscience Research; and by the McDonnell Center for Systems Neuroscience at Washington University.

