## Supplementary Materials for "The microstructure-weighted human connectome: network properties and structure-function correlations across spatial scales"

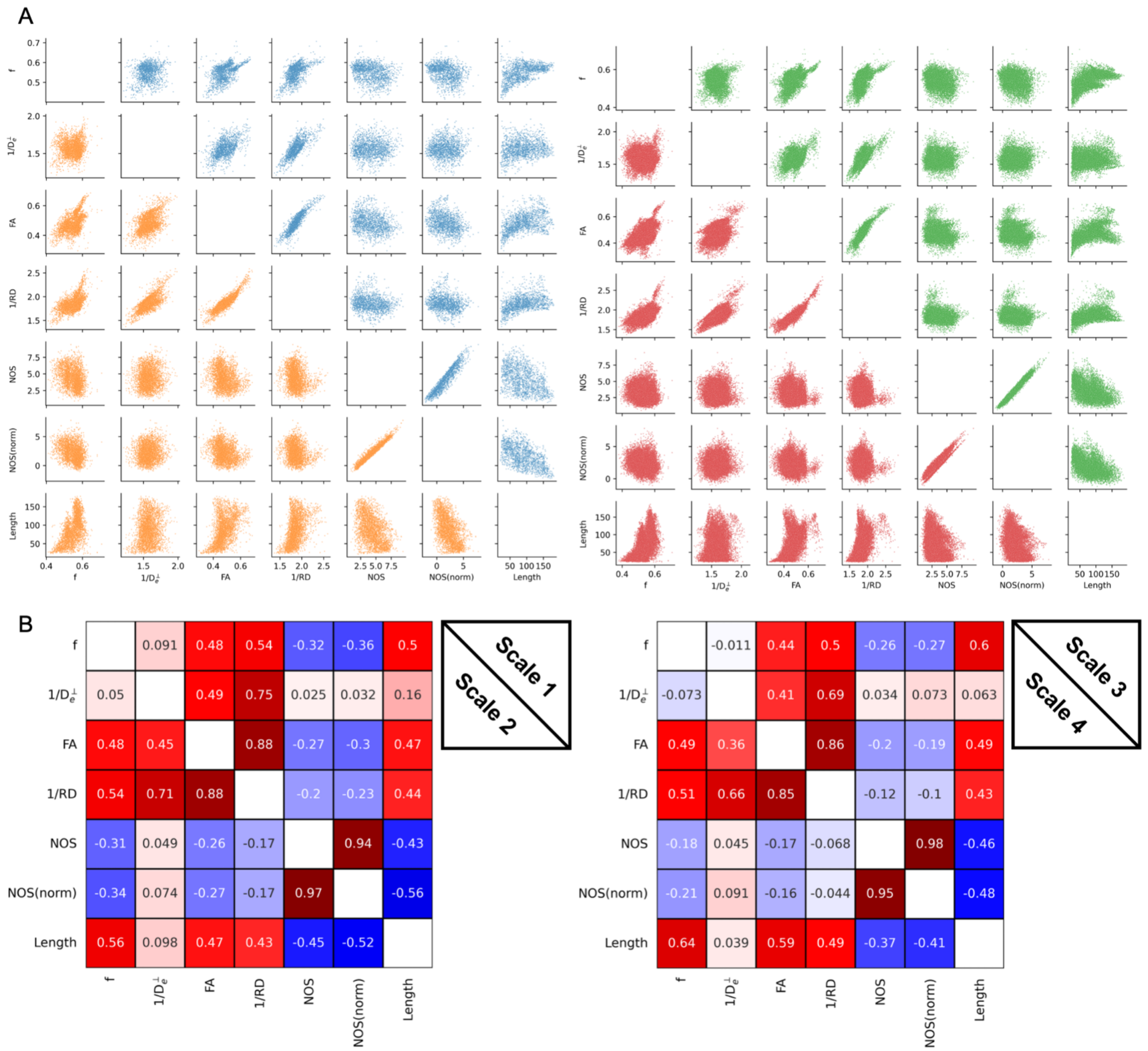


Supplementary Figure 1: Correlations between group-average connectome weights. Pairwise scatter plots (one point per edge) are shown for all parameter weights (A) with Spearman correlation coefficients (B), shown for parcellation scale 1 (left, upper triangle), scale 2 (left, lower triangle), scale 3 (right, upper triangle) and scale 4 (right, lower triangle), as indicated by the legend. Connectomes weighted by NOS and normalised NOS were log-transformed before plotting.

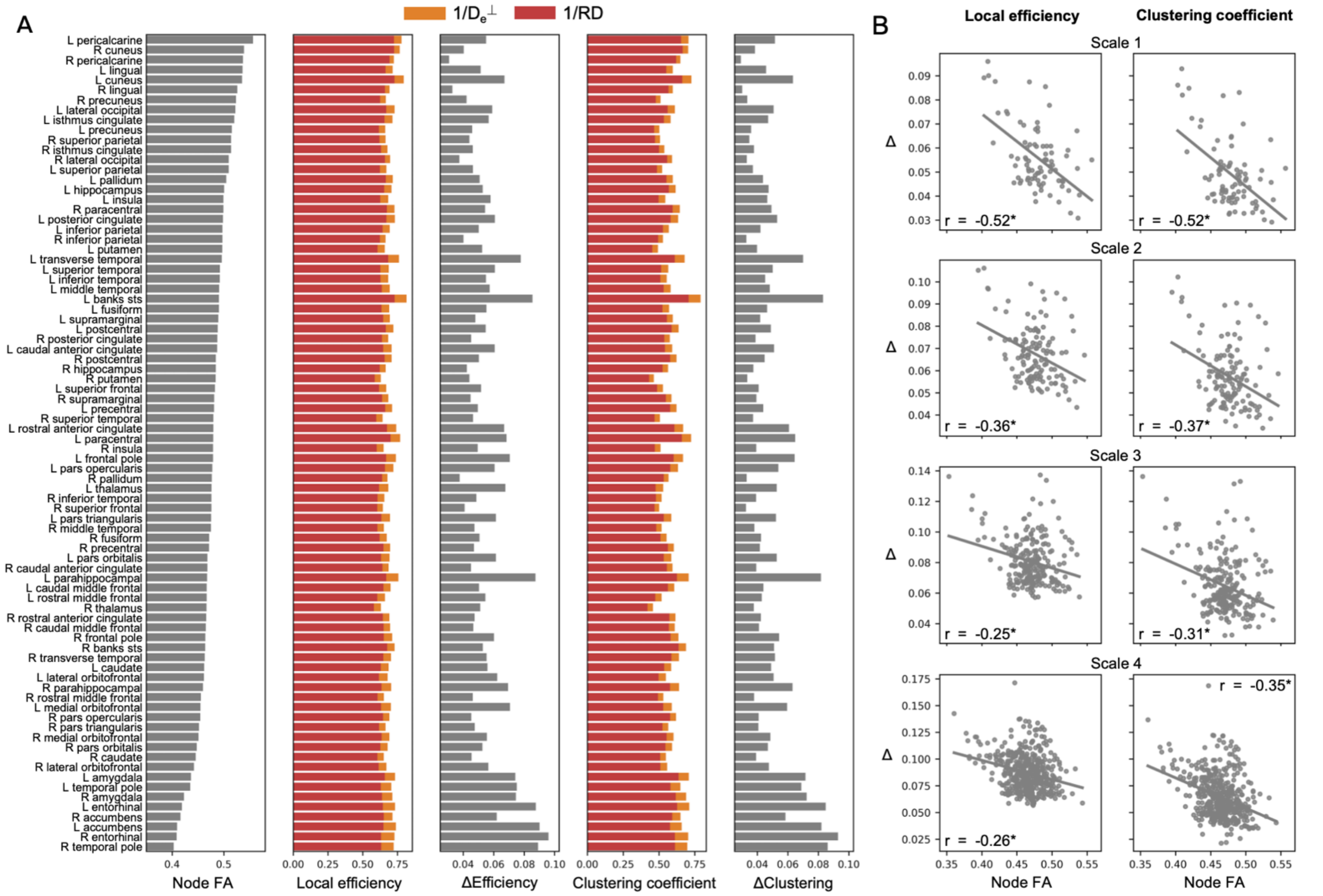


Supplementary Figure 2: Comparison of node-level measures between $1/D_e^{\perp}$ and 1/RD. Bar plots show the mean FA per node, the local efficiency for connectomes weighted by $1/D_e^{\perp}$ and 1/RD, the difference between these (Δefficiency), the clustering coefficient for connectomes weighted by $1/D_e^{\perp}$ and 1/RD, and the difference between these (Δclustering), all ordered by descending mean nodal FA, for scale 1 (A). Scatter plots show the relationship between mean nodal FA and both Δefficiency and Δclustering (B), demonstrating that the larger differences in nodal measures of network segregation between $1/D_e^{\perp}$ and 1/RD are found in the nodes with lower FA (and therefore more crossing fibres). Each scatter plot shows the Pearson correlation coefficient. *Bonferroni-corrected $p < 0.001$.

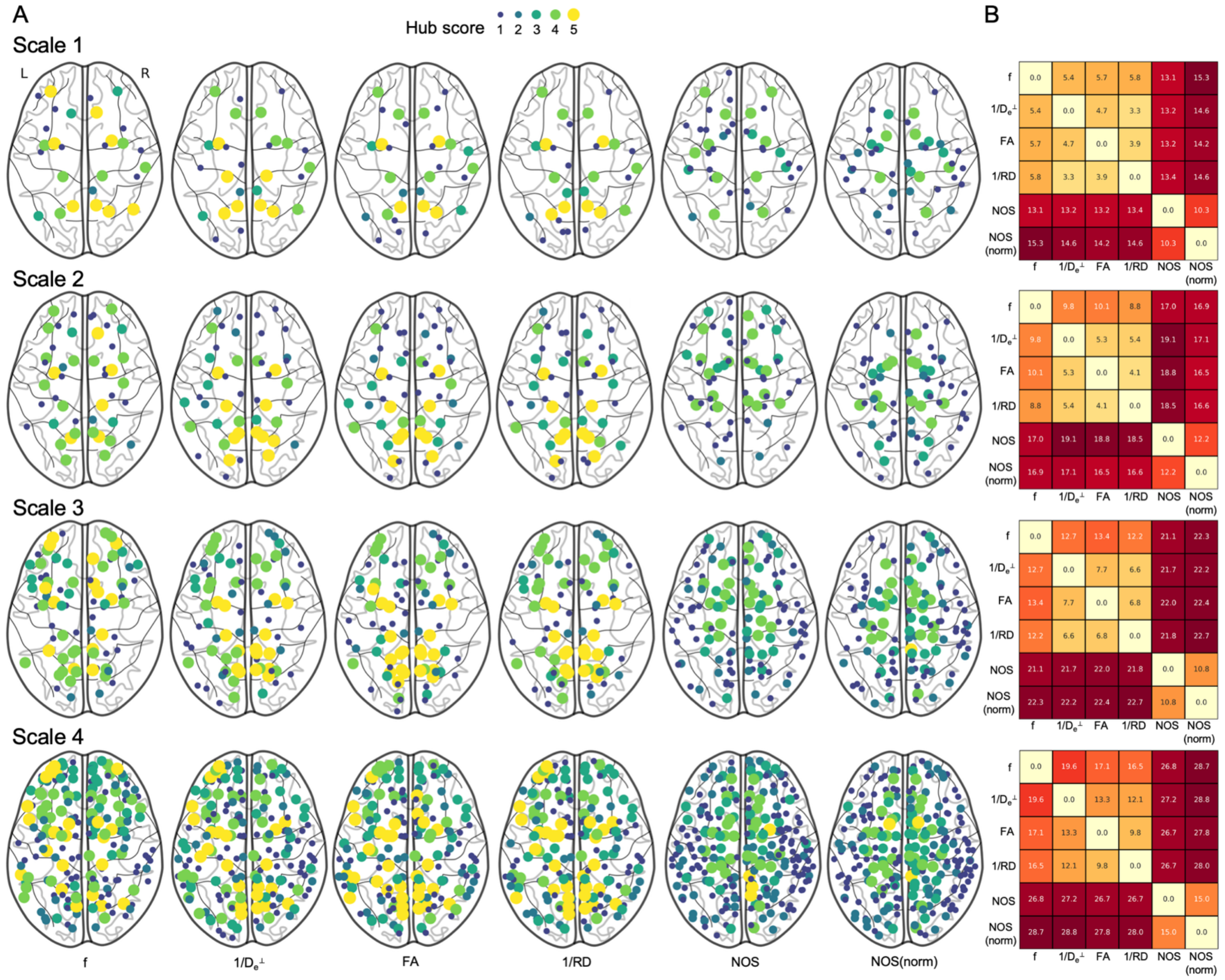


Supplementary Figure 3: Hub scores at each parcellation scale. Nodes with nonzero hub score are shown for each parameter at each parcellation scale (A), with the hub score indicated by the colour scale. The pairwise Euclidean distance between hub scores for each parameter is shown for each parcellation scale (B). Hub scores were derived by adding one to nodes in the highest 20% for node strength, betweenness, eigenvector centrality, and closeness, and in the lowest 20% for clustering coefficient.

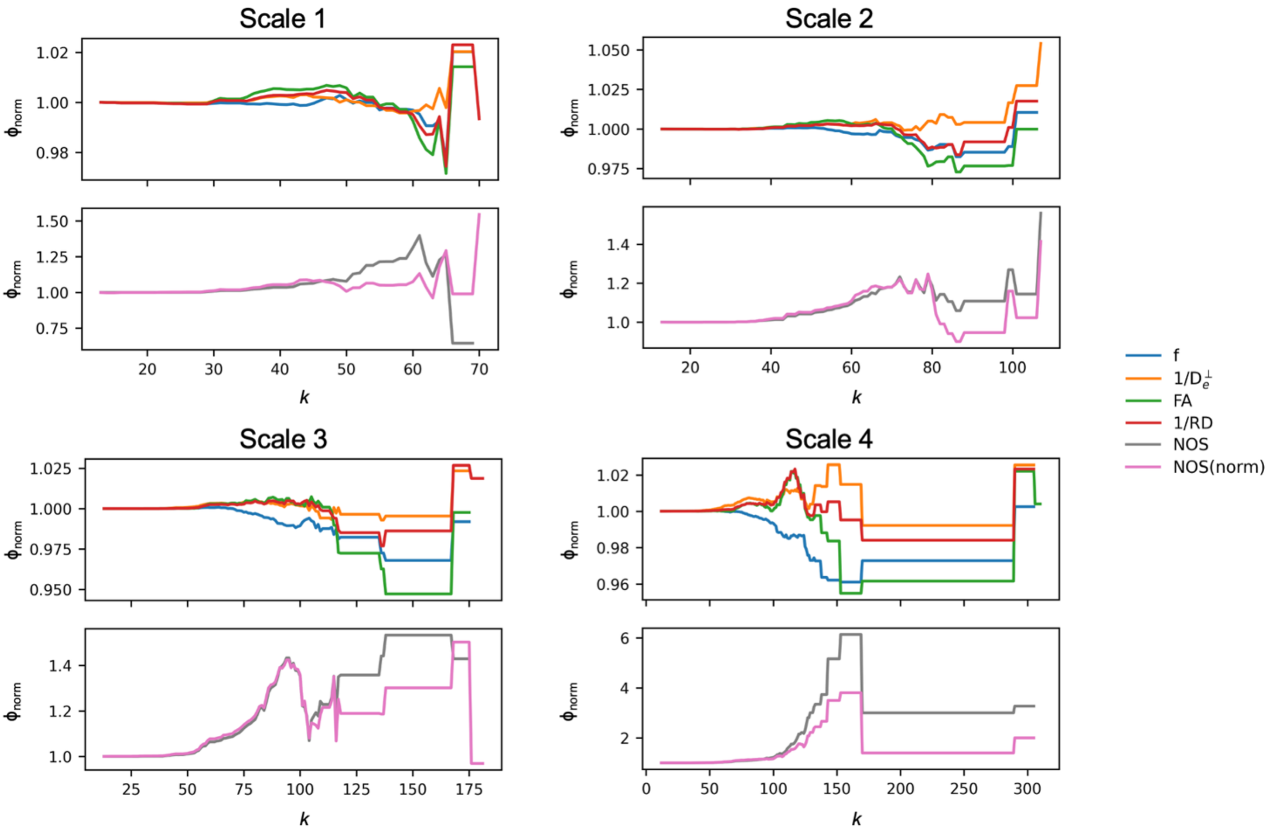

Supplementary Figure 4: Normalised rich club coefficients ($\phi_{norm}$) at each parcellation scale.

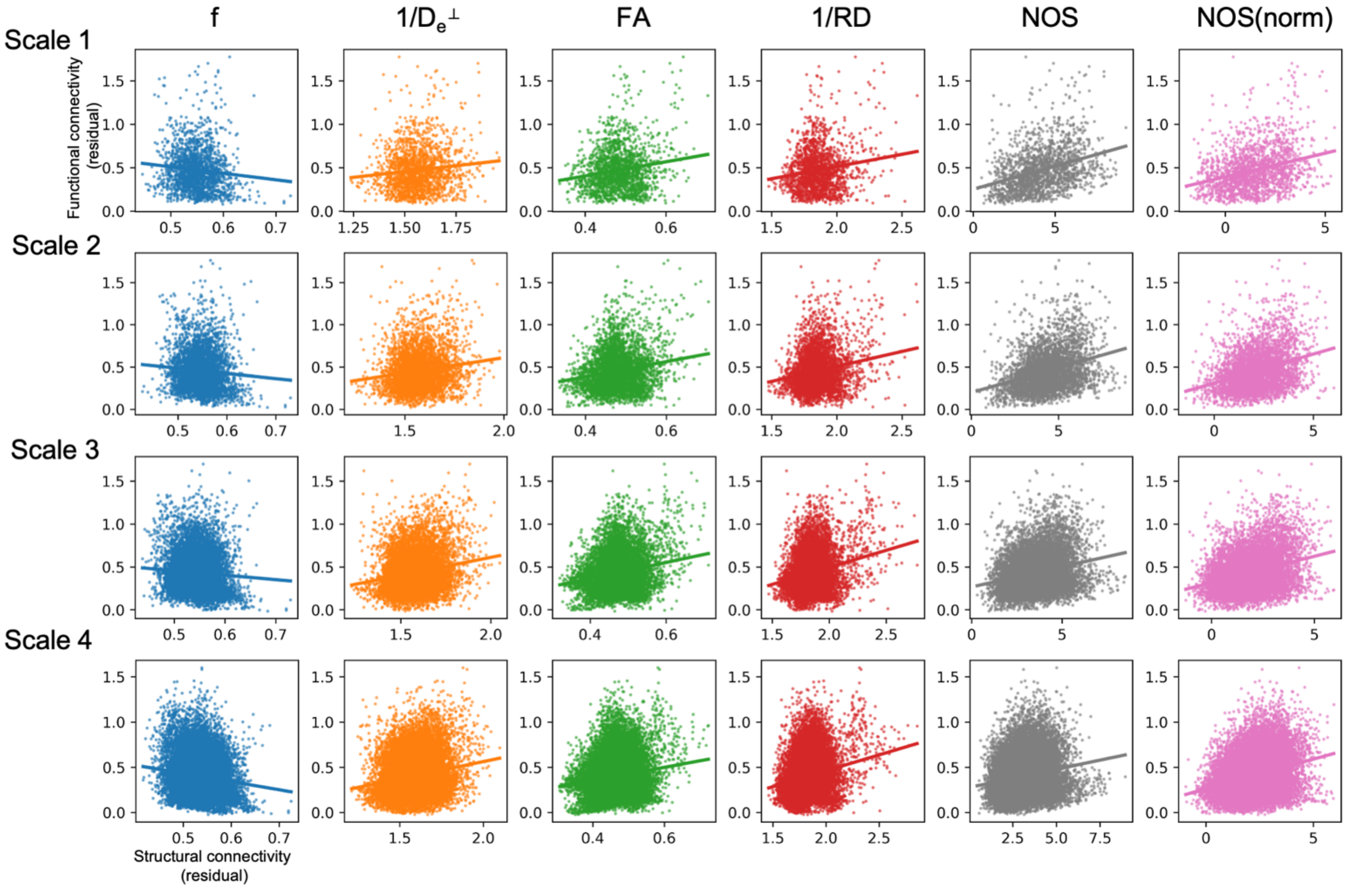


Supplementary Figure 5: Whole-brain structure-function coupling with fMRI functional connectivity. Scatter plots show the residual structural and functional connectivity after regressing length. The corresponding correlation coefficients are shown in Figure 5B in the main text.

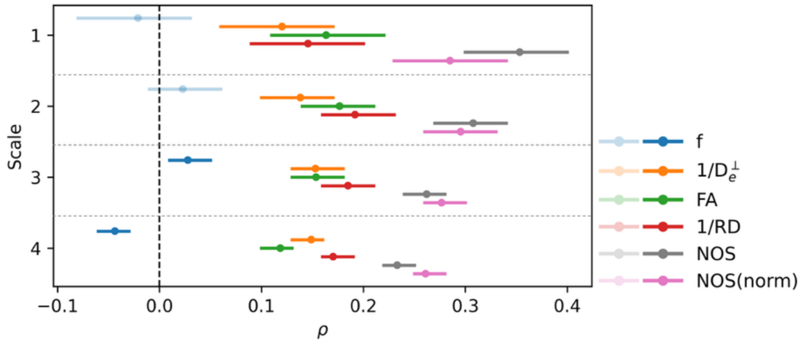


Supplementary Figure 6: Cortical structure-function coupling with fMRI functional connectivity. SFC was measured using the whole-brain approach shown in Figure 5A in the main text, but for only cortico-cortical connections (i.e. removing subcortical regions). This is plotted with bars extending to the 95% confidence intervals. Non-significant results (after FDR correction) are shown with reduced opacity.

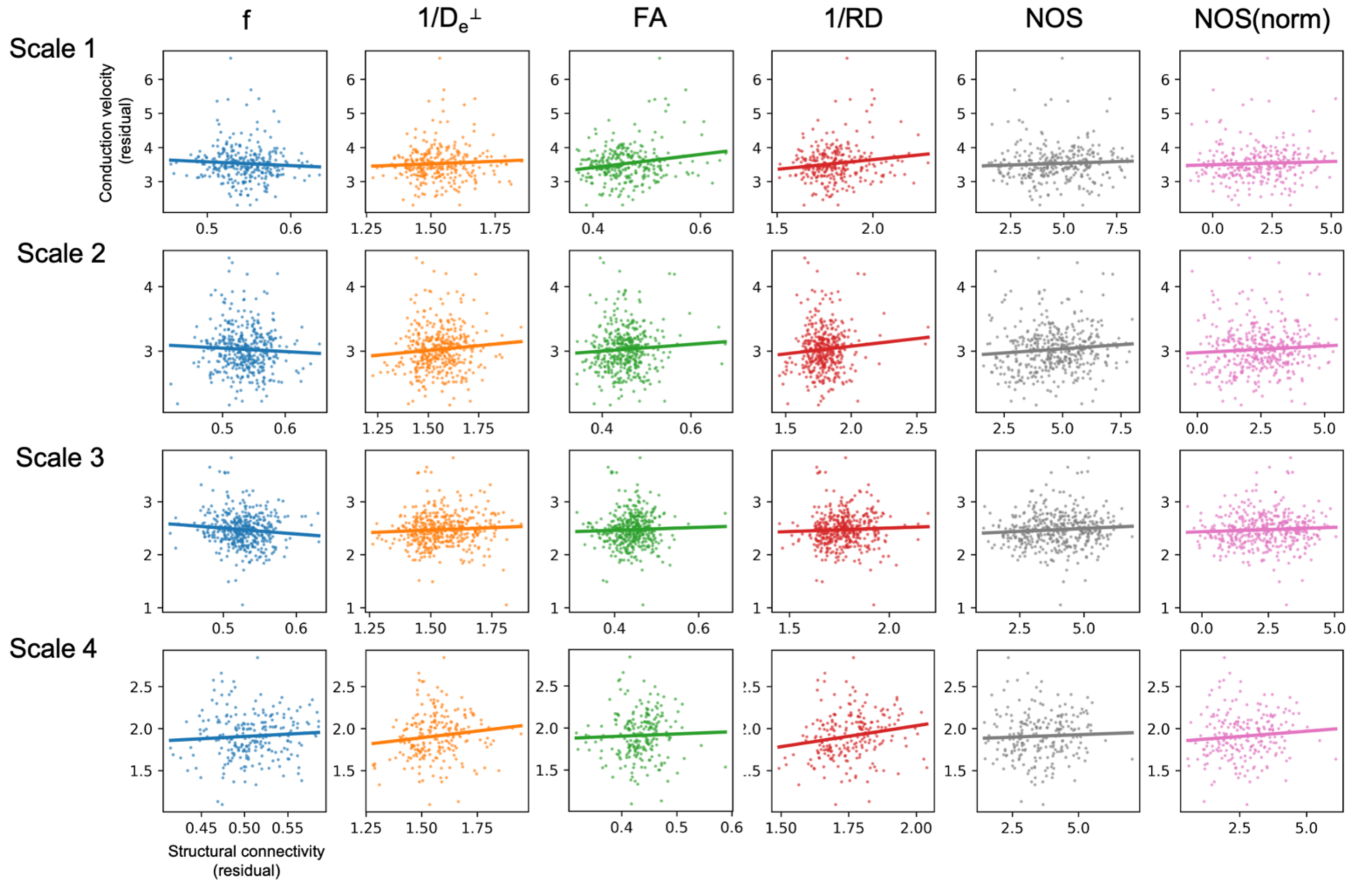


Supplementary Figure 7: Whole-brain structure-function coupling with SEEG conduction velocity. Scatter plots show the residual structural connectivity and conduction velocity after regressing length. The corresponding correlation coefficients are shown in Figure 5C in the main text.
